# The Drosophila Genome Nexus: a population genomic resource of 605 *Drosophila melanogaster* genomes, including 197 genomes from a single ancestral range population

**DOI:** 10.1101/009886

**Authors:** Justin B. Lack, Charis M. Cardeno, Marc W. Crepeau, William Taylor, Russell B. Corbett-Detig, Kristian A. Stevens, Charles H. Langley, John E. Pool

**Affiliations:** Laboratory of Genetics, University of Wisconsin-Madison, Madison, WI 53706; Department of Evolution and Ecology, University of California, Davis, CA 95616; Department of Computer Sciences, University of Wisconsin-Madison, Madison, WI 53706; Department of Integrative Biology, University of California, Berkeley, CA 94720; Department of Statistics, University of California, Berkeley, CA 94720

**Keywords:** *Drosophila melanogaster*, population genomics, genome assembly

## Abstract

Hundreds of wild-derived *D. melanogaster* genomes have been published, but rigorous comparisons across data sets are precluded by differences in alignment methodology. The most common approach to reference-based genome assembly is a single round of alignment followed by quality filtering and variant detection. We evaluated variations and extensions of this approach, and settled on an assembly strategy that utilizes two alignment programs and incorporates both SNPs and short indels to construct an updated reference for a second round of mapping prior to final variant detection. Utilizing this approach, we reassembled published *D. melanogaster* population genomic data sets (previous DPGP releases and the DGRP freeze 2.0), and added unpublished genomes from several sub-Saharan populations. Most notably, we present aligned data from phase 3 of the Drosophila Population Genomics Project (DPGP3), which provides 197 genomes from a single ancestral range population of *D. melanogaster* (from Zambia). The large sample size, high genetic diversity, and potentially simpler demographic history of the DPGP3 sample will make this a highly valuable resource for fundamental population genetic research. The complete set of assemblies described here, termed the Drosophila Genome Nexus, presently comprises 605 consistently aligned genomes, and is publicly available in multiple formats with supporting documentation and bioinformatic tools. This resource will greatly facilitate population genomic analysis in this model species by reducing the methodological differences between data sets.

## Introduction

Recent advances in next-generation sequencing have led whole genome sequencing to be extended from only a few genetic strains of select model organisms to many genomes from humans and from model and non-model taxa. While all fields of genetic analysis have benefited from these technological advances, population genetics has been especially affected as hundreds or even thousands of whole genome sequences have been generated for some organisms (e.g., The 1000 Genomes Project Consortium 2010, Pool *et al.* 2012; Huang *et al.* 2014; Long *et al.* 2013; Wallberg *et al.* 2014). As a result of these large population genomic databases, we have gained considerable power in detecting and understanding species history (e.g., Li and Durbin 2011), the genome-wide consequences of natural selection (*e.g.*, Comeron 2014), structural variation (*e.g.*, Corbett-Detig and Hartl 2012), and patterns of linkage disequilibrium and recombination (*e.g.*, Chan *et al.* 2012).

While these data sets have undeniable utility, the rapid development and deployment of next-generation technologies has been accompanied by a diversity of opinions on the most appropriate ways to assemble, filter, and curate extremely large, complex data elements. As a result, each population genomic data set is generated using unique combinations of library preparation chemistry and sequencing platform, different short-read aligning programs or pipelines with distinct biases and error rates, a wide range of quality filters and thresholds, and often distinct data formats. Ultimately, this renders population genomic data sets difficult to combine and jointly analyze. For example, it is difficult to understand whether population genetic statistics (*e.g.*, nucleotide diversity) are directly comparable given potential differences in error rate or mapping/coverage biases.

*Drosophila melanogaster* has played a pivotal role in essentially every field of genetic analysis, from population and evolutionary genetics to the development of fly models for understanding human disease. While *D. melanogaster* likely originated in Sub-Saharan Africa (Lachaise *et al.* 1988; Pool et al. 2012), natural populations now occur in essentially all temperate and tropical localities, and are typically commensal with humans. There are currently multiple independently generated population genomic data sets available that differ in the sequencing platform, assembly pipeline, and the data formats released to the public (Mackay *et al.* 2012; Huang *et al.* 2014; Langley *et al.* 2012; Pool *et al.* 2012). To address at least the last two of these issues, we present the assembly of 605 genomes from natural populations of *D. melanogaster*, all assembled using a common approach. This data set includes the previously published diploid DGRP freeze 2.0 genomes from Raleigh, NC (Huang *et al.* 2014), the DPGP collection of homozygous chromosomes from Malawi (Langley *et al.* 2012), and the haploid DPGP2 (Pool *et al.* 2012) collections of genomes from Sub-Saharan Africa. In addition, we publish 53 additional haploid embryo and inbred African genomes from Egypt, Ethiopia, Kenya, South Africa, and Uganda, along with the DPGP3 data set of 197 haploid embryo genomes from a single population in Zambia.

This deep genomic sequencing of the Zambia “ZI” sample was motivated by preliminary data suggesting that this population has maximal genetic diversity among known *D. melanogaster* populations, along with minimal levels of admixture from non-sub-Saharan (“cosmopolitan”) populations (Pool *et al.* 2012). While the sample size of DPGP3 is comparable to that of DGRP, each data set has particular strengths. DGRP includes a substantial inbreeding effort to facilitate genotype-phenotype comparisons, whereas DPGP3 uses a haploid method sequencing effort (Langley *et al.* 2011) to generate fully homozygous genomes for population genetic analysis. Because of its location within the sub-Saharan ancestral range of the species, the Zambia sample has not experienced the out-of-Africa bottleneck or New World admixture that are relevant to the DGRP population (Duchen *et al.* 2013). By providing a clear picture of diversity in the ancestral range of *D. melanogaster*, the DPGP3 collection will aid in understanding the histories of other worldwide populations and the species as a whole, as illustrated by studies of sub-Saharan human populations (CITATIONS). While Zambia may not necessarily represent a population at demographic equilibrium, the relative simplicity of its history will also facilitate studies of the effects of natural selection and other processes on genomic diversity.

The *Drosophila* Genome Nexus (DGN) created from these alignments is intended to facilitate population genetic analyses focused on single nucleotide polymorphisms (SNPs). We do not claim our assembly pipeline produces the best possible alignments. It does represent a modest advance over standard methodology, but its primary virtue is to increase the comparability of population genomic datasets. For example, there would be scientific value in comparing the North American DGRP population against separately published genomes from the European and African source populations from which it may derive (Caracristi and Schlötterer 2003; Duchen *et al.* 2013). Detailed population genetic inference is not a focus of the present study, but we present basic summaries of genetic diversity and structure, as well as patterns of admixture into sub-Saharan African populations from cosmopolitan populations.

## Materials and Methods

### Reassembly of previously published genomes

We obtained the raw sequencing reads from the NCBI short read archive (SRA; http://www.ncbi.nlm.nih.gov/sra) for Illumina data from the DGRP freeze 2.0 (Mackay *et al.* 2012; Huang *et al.* 2014), DPGP (Langley *et al.* 2012), and DPGP2 (Pool *et al.* 2012) collections of genomes (accession numbers given in Table S1). See the above citations for information concerning DNA extraction, library preparation, and sequencing, as these varied considerably among data sets.

### Newly sequenced genomes

We present a considerable expansion of the genomic sequences available for *D. melanogaster* with the addition of 246 African lines. Table S1 provides descriptions of fly stocks and their availability, genome and alignment characteristics, and raw data accession numbers. Table S2 gives information about sampling locations. This expansion consists of 193 additional lines (primarily isofemale, some inbred for five generations) collected from Siavonga, Zambia (collectively referred to as the DPGP3 data set), in addition to the 4 Zambia ZI genomes previously published (Pool *et al.* 2012). We also include here 53 additional genomes (referred to here as the African Genomes Extended Sampling, or the AGES data set) from 12 African populations. Isofemale lines for these populations were collected following Pool (2009). For all DPGP3 genomes, genomic library preparation from haploid embryos followed the protocol of Langley *et al.* (2011). Sequencing for DPGP3 was performed on an Illumina Genome Analyzer IIx (Langley lab, UC Davis). From the AGES data set, all but the Egyptian (EG) and Kenyan (KM) paired-end libraries were prepared from haploid embryos using the same methods as for DPGP3. All AGES genomes were sequenced at the UW-Madison Biotechnology Center on the Illumina HiSeq 2000 platform. For the three EG genomes, inbred lines were established through full-sib mating for 8 generations, while the four KM genomes were sequenced directly from isofemale lines that had been maintained in the laboratory for 12 years and therefore passively inbred. For these 7 genomes, we extracted DNA from 30 females, paired-end library preparation and size-selection for 300 bp inserts was conducted using the NEBNext DNA Library Prep Reagent Set (New England BioLabs), and sequencing was conducted at the UW-Madison Biotechnology Center on the Illumina HiSeq 2000 platform.

### Genome assembly pipeline

In reference assembly of short-read sequencing data, a major limitation is SNP and indel divergence of sequenced genomes from the reference. This issue could result in alignment biases if reads derived from low diversity regions of the genome can be aligned more confidently, in addition to biases among genomes that vary in their overall divergence from the reference sequence (*e.g.* sub-Saharan vs. cosmopolitan *D. melanogaster*). In an attempt to ameliorate such effects, we developed and applied a pipeline that combines two aligners with different degrees of sensitivity to non-reference variation and speed, and utilizes two rounds of mapping (Fig. S1). In brief, we first mapped short read data to the *D. melanogaster* reference genome (release 5.57; http://flybase.org) using BWA v0.5.9 (Li and Durbin 2010) using default settings, followed by mapping of all unmapped reads using Stampy v1.0.20 (Lunter and Goodson 2010). This approach combines the rapid but strict BWA algorithm to first map the relatively “easy-to-align” reads, with the more sensitive but computationally intensive Stampy algorithm, which more effectively and accurately aligns the relatively divergent reads (Lunter and Goodson 2010). All reads with mapping quality scores below 20 were excluded. Optical duplicates were then removed using Picard v1.79 (http://picard.sourceforge.net/) and assemblies were improved around indels using the GATK v3.2 Indel Realigner (cKenna *et al.* 2010; DePristo *et al.* 2011). The Unified Genotyper (DePristo *et al.* 2011) was then used to call indels and SNPs for each individual genome. Among the indel calling criteria, >50% of the reads at a given position had to support the existence of that indel, with a minimum of 3 reads containing the variant. For SNP calling in this first round, we required a minimum phred-scaled quality value of 31, and that >75% of reads at a given position support the SNP. For the second round of assembly, the SNPs and indels called in the first round were introduced into the *D. melanogaster* reference, and this modified reference was then used for a second round of mapping. Following indel realignment, the Unified Genotyper was then used to call all sites in the modified reference genome. To generate reference-numbered consensus sequences, a custom perl script was used to shift all base coordinates back to those of the original *D. melanogaster* reference. Deletions and all sites within 3 bp of a called indel were coded as “N” (based on the error analysis described in the results section), while insertions do not appear in reference-numbered consensus sequences.

### Consensus error rate and sequence generation

To estimate the actual error rate of our assemblies, and to determine the optimal trade-off between error rate and genomic coverage (the number of euchromatic bases with called alleles), we evaluated base-calling accuracy using the previously published resequenced reference genome (*y*^1^ *cn*^1^ *bw*^1^ *sp*^1^; Pool *et al.* 2012), sequenced on a GAIIx to ∼25X average depth with 76 bp paired end reads (Table S1). Variation was simulated via dwgsim (https://github.com/nh13/DWGSIM/wiki); we introduced substitutions randomly across the genome at a rate of 0.012/bp, with an indel rate of 0.002/bp (with a probability of 0.6 of indel extension). This variation was used to produce a modified reference sequence, as in Pool *et al.* (2012). The resequenced reference reads were then mapped to the modified reference using the pipeline described above, as well as several variations, to investigate the performance of our pipeline versus more standard alignment approaches and various degrees of filtering. Analysis of simulated sequence reads via dwgsim gave highly concordant results (not presented).

### Heterozygosity filtering

For the DGRP data set, the EG and KM samples from the AGES data set, and the ZK genomes from the DPGP2 data set, libraries were constructed from pools of flies following varying degrees of inbreeding. For these genomes, tracts of heterozygosity can remain (and can even be substantial), presumably due to the presence of multiple recessive lethal and sterile mutations that are segregating in repulsion. These linked lethals may often occur, for example, within large inversions that are polymorphic within a line and suppress crossing over. To allow consistency between haploid and diploid genomes, entire heterozygous regions must be filtered out prior to generating homozygous consensus sequences.

For the samples mentioned above, the Unified Genotyper was run in diploid mode to enable the calling of heterozygous sites. To identify and mask residually heterozygous regions, we scanned the 5 euchromatic arms of each diploid genome for heterozygous calls in 100 kb windows advancing in 5 kb increments. Rather than use a hard boundary for delineating windows of residual heterozygosity, we chose to scale the threshold for a given window to the level of genetic diversity observed in that window within either sub-Saharan or cosmopolitan populations (henceforth referred to as *π*_*sub*_ and *π*_*cos*_, respectively), depending on the geographic origin of each individual genome. To determine these thresholds, we estimated nucleotide diversity (*π*) in 100 kb windows advancing in 5 kb increments for the large Rwandan (RG) sample of 27 haploid embryo genomes and the French sample of 9 haploid embryo genomes to represent sub-Saharan and cosmopolitan diversity, respectively. If the proportion of heterozygous sites in a given window exceeded *π*/5, that window was enucleated, and this window was extended in both directions on the chromosome arm until encountering a window with heterozygosity less than *π*/20. This procedure was conducted beginning from both ends of each chromosome arm. Tracts of heterozygosity were masked to N, and are provided in Table S3.

For the DGRP data set, a subset of these genomes had elevated baseline levels of heterozygosity for unknown technical reasons. For the majority of these genomes, this constituted <10% of all sites, while 29 genomes had >10% of sites masked for this reason (Table S3). Regions with elevated numbers of putatively heterozygous sites were masked from consensus sequences regardless of whether they reflected true heterozygosity, cryptic structural variation, or technical artefacts. However, we also used a normalization approach to identify the DGRP genome regions that reflect genuine heterozygosity. We generated normalization factors using the following procedure: (1) for each euchromatic chromosome arm in each genome, first determine the mode of heterozygous calls per site (hets/site) in the same windows above (in bins of 0.00001), only including windows with a hets/site between 0 and that window’s *π*_*cos*_/2 to remove the effects of true heterozygosity in determining the mode of the baseline (“genomic noise factor”); (2) obtain each genome’s normalization factor by dividing the above genomic noise factor by the mode of all DGRP genomic noise factors, truncating this normalization factor at 1 (since we are only interested in reducing the influence of non-genuine heterozygosity calls). (3) appropriate for the identification of true heterozygosity, using the criteria described above.

While haploid embryo genomes are not expected to contain any true heterozygosity, repetitive and/or duplicated regions can cause mismapping that results in tracts of “pseudoheterozygosity”. To detect these tracts and remove them, we implemented the same threshold approach as outlined above (without normalization, since none of these genomes showed elevated background levels of putative heterozygosity). For these genomes, the Unified Genotyper was run in haploid mode, and so read proportions were analyzed in place of called heterozygous sites. For windows with the proportion of sites with <75% of the reads matching the consensus base above *π*/5, that window was enucleated, and this window was extended in both directions until encountering a window below *π*/20. This procedure was conducted starting from both ends of the chromosome arms, overlapping windows were merged, and all pseudoheterozygosity tracts are reported in Table S3.

### Chromosomal inversion detection

Chromosomal inversions are known to be common in natural *D. melanogaster* populations (*e.g.*, Krimbas and Powell 1992; Aulard *et al.* 2002), and can significantly impact the distribution of genetic diversity (*e.g.*, Kirkpatrick and Barton 2006; Hoffman and Rieseberg 2008; Corbett-Detig and Hartl 2012). For the Drosophila Genome Nexus, we compiled known inversions for the previously published genomes and also identified inversions for the newly sequenced genomes. For DGRP genomes, inversions were previously identified cytogenetically (Huang *et al.* 2014). For the DPGP2 dataset, common inversions were previously detected using the approach of Corbett-Detig *et al.* (2012). For the newly sequenced DPGP3 and AGES datasets, inversions were also detected using this method, and we provide the identified inversions for all of the analyzed genomes in Table S5.

### Detection of identical-by-descent (IBD) genomic regions

Tracts of IBD may reflect the sampling of related individuals, and can contradict theoretical assumptions and complicate many population genetic analyses. To identify tracts of IBD, we implemented the approach of Pool *et al.* (2012), but with slight modifications for the diploid genomes and for the large DPGP3 population sample (described below). All possible pairwise comparisons were made for each of the five euchromatic arms of each genome, and pairwise differences per site were calculated in 500 kb windows advanced in 100 kb increments. Windows with less than 0.0005 pairwise differences per site were deemed putatively IBD. Some chromosomal intervals (including centromere- and telomere-proximal regions) exhibited large-scale, recurrent IBD between populations suggesting explanations other than close relatedness, and therefore did not contribute to a genome’s IBD total unless they extended outside these recurrent IBD regions. Elsewhere, within population IBD (presumably due to very recent common ancestry) was determined to be that which totaled genome-wide >5 Mb for a pairwise comparison of genomes.

For the DPGP2 and AGES genomes, we excluded the same recurrent IBD regions as those of Pool *et al.* (2012). However, for the much larger DGRP and DPGP3 samples of genomes, we visually reexamined these recurrent IBD tracts and generated new regions to be excluded for each of these data sets (provided in Table S4).

Due to heterozygosity filtering, some diploid genomes had genomic coverages far less than the typical ∼111 Mb. Therefore, the genome-wide threshold for IBD filtering was adjusted to 5% of all called positions rather than 5 Mb. In addition, only 500 kb windows with >100 kb pairwise comparisons were allowed to contribute to the 5% total, minimizing the influence of windows with large numbers of masked sites.

### Detection of cosmopolitan admixture

Because admixture from cosmopolitan gene flow into Africa can significantly impact estimates of genetic diversity and violate demographic assumptions of some analyses, it is important to identify instances of cosmopolitan ancestry in the sub-Saharan genomes. We used the HMM approach outlined in detail by Pool *et al.* (2012), but with updated reference panels. The sub-Saharan reference panel included 27 Rwanda (RG) genomes, but chromosome arms with known inversions were excluded (as identified by Corbett-Detig and Hartl 2012). The cosmopolitan reference panel included 9 France genomes, again excluding inversions, since inverted arms were previously found to have unusually high divergence from standard arms in this population (Pool *et al.* 2012; Corbett-Detig and Hartl 2012). Pairwise distance comparisons indicated that the Egyptian genomes were genetically cosmopolitan. To augment the cosmopolitan sample, we included homozygous regions of standard arrangement Egypt chromosome arms in the cosmopolitan reference panel.

Aside from these modifications to the reference panels, we implemented the admixture HMM as described in Pool *et al.* (2012). Briefly, this HMM works in the following way. For a focal African genome within a particular window, the method compares it against a cosmopolitan reference panel and assesses whether its genetic distance to this reference panel is on the level expected for a sub-Saharan genome, or if instead this resembles a comparison of one cosmopolitan genome against others (indicating admixture). As before, window size for the analysis was based on 1000 non-singleton SNPs among the RG sample, roughly corresponding to a mean window size of 50 kb. The analysis was initially calibrated using the 27 RG genomes to represent the putative non-admixed state, and emission distributions for the non-admixed state were generated as in Pool *et al.* (2012). A revised sub-Saharan panel was then generated through an iterative analysis of the RG genomes. Following a single round of the method, RG genomes were masked for admixture, and then these masked genomes served as the African panel for a second round. RG genomes were then masked for admixture again, and a third round of the method was applied to the RG genomes to produce a final set of emissions distributions that were used in the analysis of all other African genomes.

### Genetic diversity and population structure

For all of the analyses described below, only heterozygosity- and IBD-filtered genomes were utilized. Sub-Saharan genomes were also filtered for cosmopolitan admixture as detailed above. Nucleotide diversity (*π*) was initially calculated in windows of 2,000 non-singleton RG SNPs, corresponding to a median window size of 100 kb for all populations with at least two genomes sampled. For more efficient analysis of the large Raleigh (DGRP) and Siavonga, Zambia (DPGP3) populations, we selected 30 genomes from each of these populations with the highest genomic coverage and with at least 30X average depth. To remove the effects of spurious estimates due to low coverage windows, we excluded windows for a given population if site coverage (the number of sites with alleles called for two or more genomes) was below half the coverage in the large RG sample for that window. To obtain whole-arm and genome-wide estimates, we conducted a weighted average of windows (weighted by the number of sites in each window with data from at least two genomes).

To examine patterns of population structure, we calculated *D*_*xy*_ and *F*_*ST*_ (Hudson *et al.* (1992) for all populations with at least two high-coverage genomes (after IBD and admixture filtering), and including the *D. melanogaster* reference genome for *Dxy*. Both analyses were conducted in windows of 2,000 non-singleton RG SNPs, and a weighted average of windows was used to obtain whole-arm and genome-wide estimates. In addition, to lessen the influence of large inversions on estimates of genetic diversity and population structure, we estimated nucleotide diversity and pairwise *F*_*ST*_ excluding inverted arms for a subset of populations with larger sample sizes (inversion presence/absence is given in Table S5).

## Results

### The Drosophila Genome Nexus

The resulting data set, which we have named the Drosophila Genome Nexus (http://www.johnpool.net/genomes.html), consists of 605 sequenced genomes (varying slightly in number among the 5 euchromatic chromosomal arms) from 36 populations from Africa, Europe, and North America. The consensus sequences analyzed below and made available online include only the 5 euchromatic chromosome arms. These consensus sequences have been filtered for heterozygosity, with additional files provided to facilitate masking of IBD and cosmopolitan admixture as well as locus-specific analysis. SNP and indel variant call files (VCFs) are also available online, both for these 5 arms and for other arms (mitochondria, chromosomes 4 and Y, and heterochromatic components of the euchromatic arms). The repetitive nature of non-euchromatic arms may entail much higher error rates; we do not focus on their analysis here.

While the consensus sequences made available online specifically focus only on SNP variants, the provided indel vcf files will also be of considerable utility. For indels, the Unified Genotyper is limited to detecting only those encapsulated entirely within a single read. Therefore, read lengths will limit the size of detected indels. To examine the extent of this effect on indel detection, we examined indel length distribution for two DPGP3 genomes with 76 bp paired-end reads as well as for two DPGP3 genomes with much longer read lengths (146 and 150 bp; Fig. S2), each with similarly low cosmopolitan admixture and high mean depth. For indels approximately 25 bp or shorter, the long and short read lengths appeared to have no effect on indel length frequencies. However, for longer indels (>25 bp) the gap in detection between the two read lengths gradually increased, illustrating the decreasing ability of the present approach to detect indels as they approach the read length. This potential bias is important to consider when examining the provided indel calls. A more comprehensive analysis of structural variation within and between these populations will be a target of future research.

While there is considerable variation among genomes in terms of average sequencing depth and coverage, the majority of this variance lies in the DGRP and AGES data sets, which range from approximately 12X to more than 100X mean depth, while the remaining genomes are primarily haploid embryo genomes of roughly 30X mean depth or higher (Table S1). In addition, coverage varies considerably among the inbred/isofemale genomes from the AGES and DGRP data sets due to heterozygosity filtering.

### Genome assembly pipeline performance

To investigate pipeline performance, basecalling bias, and consensus error rate, we assembled a resequenced *D. melanogaster* reference strain to an artificially mutated reference genome to simulate variation. Overall, adding a second round of mapping that incorporated SNP and indel variants called in the first round of mapping resulted in approximately a 1% increase in sequence coverage (just over 1 million sites added) and a significant improvement in error rate relative to performing only a single round of mapping with only BWA (Fig. 1). This improvement was observed irrespective of the nominal quality value threshold for basecalling (which had only a modest effect on error rates), with error rates for the two round assemblies completely distinct from the distribution of error rates observed for a single round of assembly. We investigated the contribution of Stampy to this improvement and found that, while error rate and coverage both improved, the vast majority of improvement was due to adding the second round of mapping (Fig. 1).

**Figure 1.**
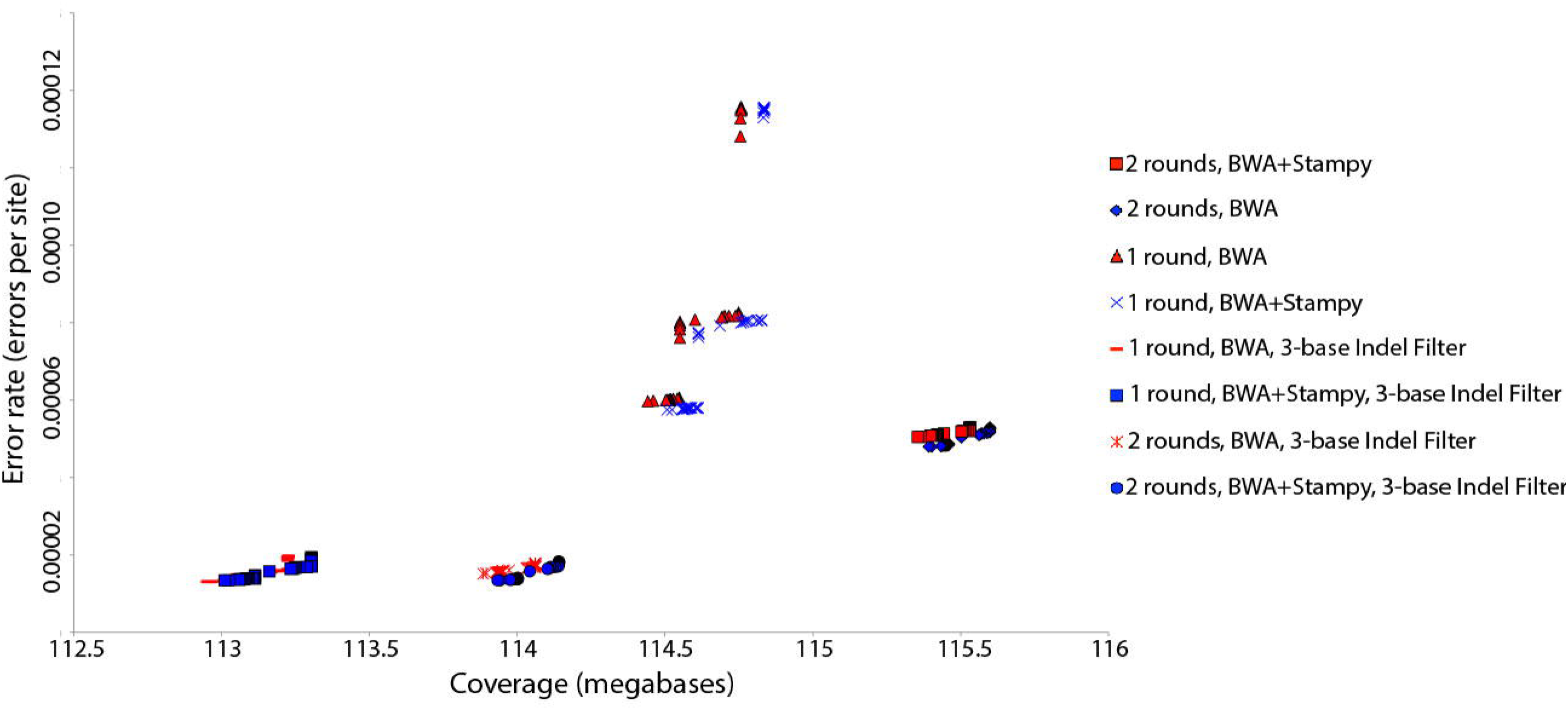
Comparison of genomic coverage and error rate for several genome assembly pipeline variations, based on resequencing of the *D. melanogaster* reference strain (Pool *et al.* 2012). All quality values from Q10 to Q100 are shown; many gave very similar results.

We also investigated the impact of filtering around indels, as past analyses have found that positions directly adjacent to indels are difficult for aligners to correctly align and a major contributor to error (Meader *et al.* 2010; Alkan *et al.* 2011). We found similar results, with approximately a five-fold reduction in error rate by masking 3 basepairs on either side of consensus indels. Assessing the possible benefit of masking 5 bases rather than 3, we observed almost no improvement in error rate to justify the nearly 1% reduction in coverage, and therefore used the 3 bp mask. Our use of the GATK Indel Realigner (McKenna *et al.* 2010; DePristo *et al.* 2011), in conjunction with incorporating indels into the reference used in the second round of mapping, may have improved our ability to align around these regions. Finally, to determine the optimal alignment quality value threshold for consensus sequence generation and to estimate the expected error rate of our assemblies, we examined the tradeoff between coverage and error rate at a range of quality values for both the haploid and diploid callers of the Unified Genotyper (DePristo *et al.* 2011), and we selected a minimum of Q75 and Q32 for calling a position in haploid or diploid genomes, respectively (Figs. S3 and S4, respectively). These thresholds corresponded to an error rate of roughly 1.36 × 10^−5^ errors per site.

To further examine our two round pipeline performance, we compared sites called only by the two round pipeline versus those called using just a single round of BWA to map, and estimated both error rate and diversity for both classes of sites. In terms of error rate, sites called in both pipelines possessed an error rate of 9.3 × 10^−6^, just below the genome-wide rate, while sites added only in our two round pipeline had an error rate of 2.9 × 10^−5^, roughly two fold higher than our genome-wide average. This increase in error rate is not surprising given that these sites added by our two round pipeline likely constitute highly diverse, hard-to-align regions relative to those confidently called by both pipelines.

To further examine the sites added by our two round pipeline, we identified bases called in both pipelines vs. those called only in our two round pipeline for a single RG genome (RG33), and then calculated nucleotide diversity at each class of sites for the RG sample of 27 genomes. In addition, for two RG genomes sequenced at similar depths (RG33, RG5) we determined both the number of sites added by our two round pipeline and the number of indels (indel “rate”) in 100 kb non-overlapping windows. For sites called only by our two round pipeline, nucleotide diversity was over three fold higher than that for sites called in both pipelines (Table 1), and we observed a clear positive relationship between the number of sites added with the new pipeline and the number of indels in that region of the genome (Fig. 2). These lines of evidence support the idea that the sites added by our pipeline are found in high diversity, difficult-to-align genomic regions.

**Table 1.**
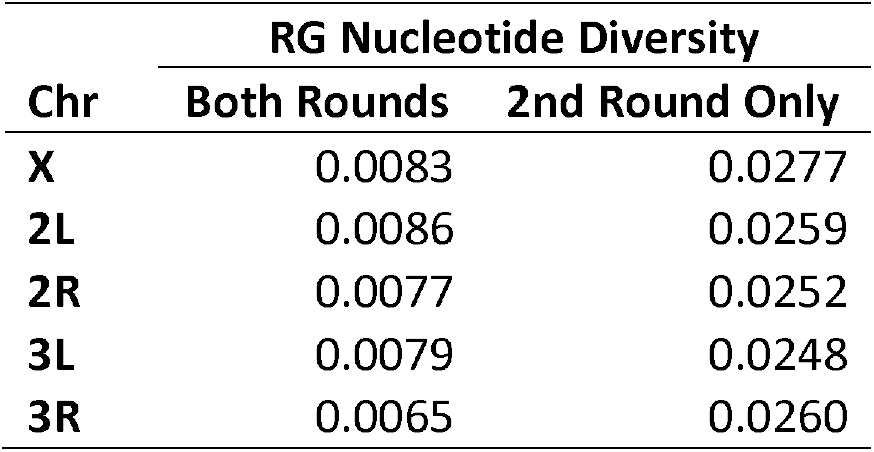
Chromosome arm nucleotide diversity (*π*) for the RG population based on sites called in both rounds of our pipeline, and for sites called only by adding the second round of mapping.

| Chr | RG Nucleotide Diversity |  |
| --- | --- | --- |
|  | Both Rounds | 2nd Round Only |
| **X** | 0.0083 | 0.0277 |
| **2L** | 0.0086 | 0.0259 |
| **2R** | 0.0077 | 0.0252 |
| **3L** | 0.0079 | 0.0248 |
| **3R** | 0.0065 | 0.0260 |

**Figure 2.**
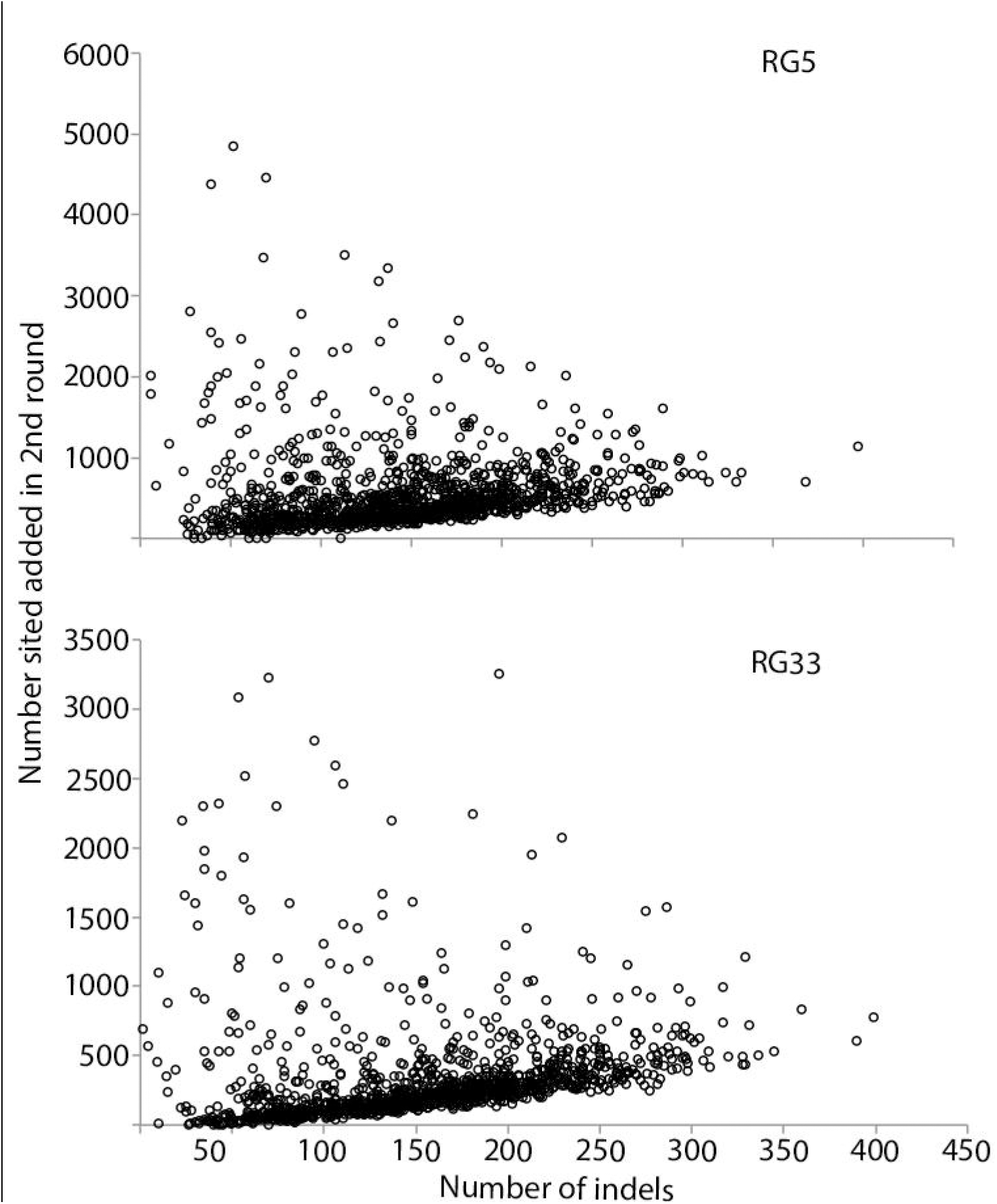
Relationship between the number of indels and the number of sites called by our two round pipeline but not called in a single-round pipeline for two RG genomes (RG5 and RG33). Site counts (y axis) and indel counts (x axis) were determined in 100 kb windows across each genome.

To determine whether any particular functional class of sites contributed disproportionately to the sites added by our two round pipeline, we used the *D. melanogaster* reference genome annotations (v5.57) to assign each individual site called only by our two round pipeline for RG33 and RG5 to one of 9 site classes: Nonsynonymous, 2-, 3-, or 4-fold synonymous, 5’ UTR, 3’ UTR, intronic, short intron (Halligan and Keightley 2006), or intergenic. While all classes of sites contributed to the total sites added by our two round pipeline, only intergenic and intronic sites were positively enriched for both RG genomes (Fig. 3), suggesting our pipeline disproportionately added these two functional classes of sites to the assemblies. However, the representation of each functional category in the “added sites” class is fairly close to null expectations, and we even added 41,368 and 36,990 nonsynonymous sites, which we would expect to be the least diverse and therefore easiest to align confidently, to RG33 and RG5, respectively, by applying the full pipeline. We also characterized the tract length and genomic distribution of sites added in the second round of our assembly for both RG5 and RG33. In terms of tract length, the vast majority of bases added occurred in short tracts of 1 to 10 bases (Fig. S5). To examine genomic location of these sites, we calculated the number of sites in 100 kb windows across the 5 euchromatic arms of the genome (Fig. S6). While sites were added somewhat uniformly across the genome, repetitive telomeric regions were especially enriched.

**Figure 3.**
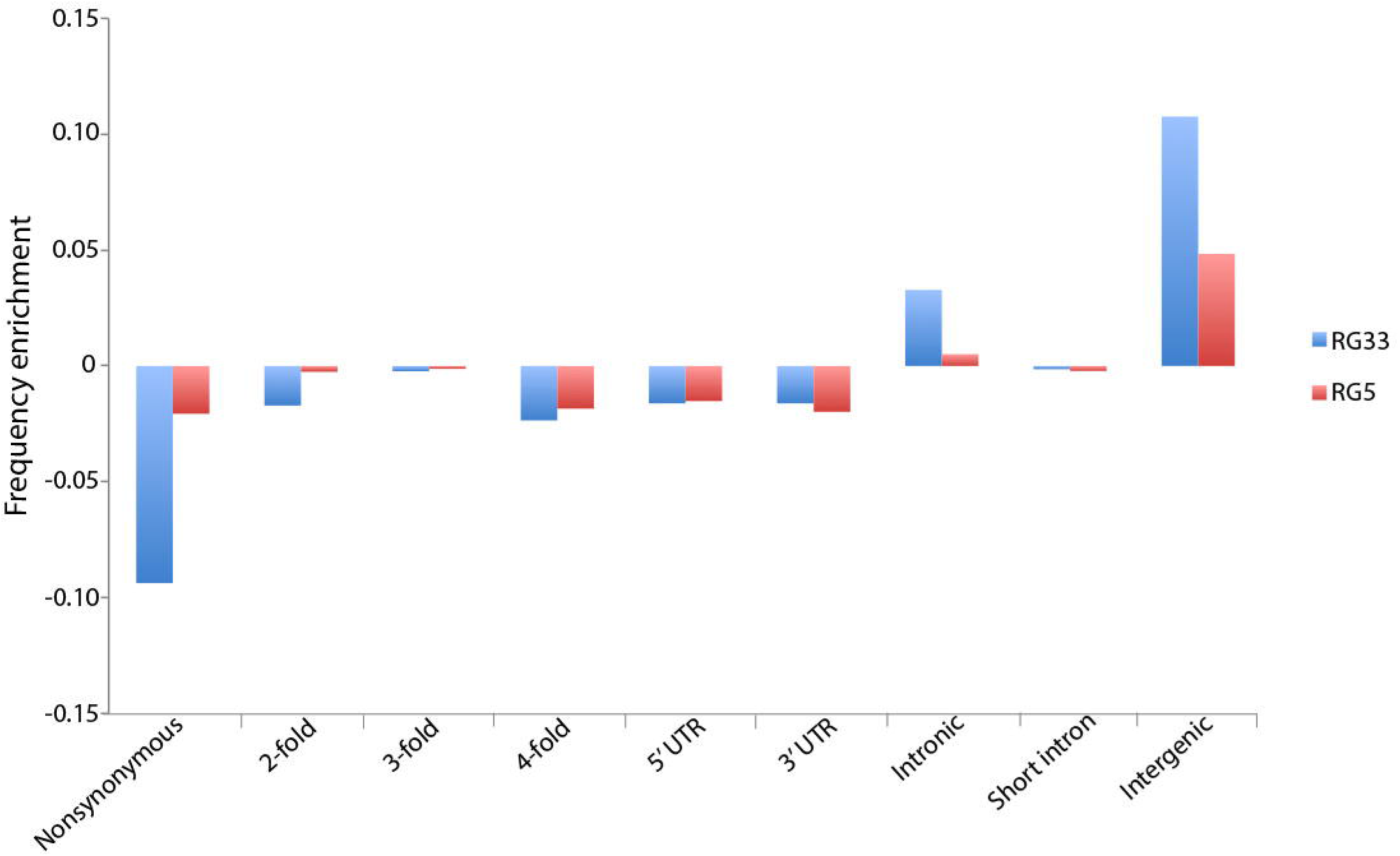
Enrichment of each of 9 annotation classes in the sites added by our two round pipeline, but not called by a single round pipeline, relative to genome-wide frequencies. We examined two RG genomes (RG5 and RG33) with approximately 30X mean depth and comparable coverage.

### Impact of sequencing depth on genetic distance

In a previous assembly of the DPGP2 data set, Pool *et al.* (2012) found a positive, non-linear relationship between mean sequencing depth (the average number of reads per basepair) and genetic distance to both the reference and the Siavonga, Zambia (ZI) population. This relationship is especially pronounced below 25X mean depth. Here, we ameliorated that issue by using a consensus caller that is less vulnerable to reference sequence bias and by adding more stringent quality filtering (http://www.dpgp.org/dpgp2/DPGP2.html). To examine the impact of this basecalling bias in our pipeline, we quantified the recall rate of reference and non-reference alleles in the resequenced reference. The recall rate for reference and non-reference alleles was nearly identical (0.958 vs 0.959), suggesting reference bias has a minimal effect. To further examine this relationship for our two round pipeline versus the single round of mapping with only BWA (but including the indel filter), we calculated mean genome-wide distance to the ZI population for each of the genomes in the AGES data set (excluding genomes with whole arms masked due to heterozygosity). For the single round of mapping, a positive relationship between mean depth and distance to ZI was apparent below 20X mean depth, but for our two round pipeline distances remained flat even approaching 10X depth (Fig. 4A). When limiting this analysis to sites called in all analyzed genomes, the bias observed below 20X mean depth for single round genomes disappeared, and distance estimates were essentially identical to those of the two round pipeline (though greatly reduced in both cases, reflecting the exclusion of more diverse genomic regions). These results suggest that the depth-related bias observed for the single round alignments (Fig. 4A) was not due to biased consensus-calling (since that would still affect the filtered analysis), but instead stems from differences in genomic coverage between low and high depth genomes. And indeed, we observe that genomic coverage is more dependent on depth in the single round alignments than for the full pipeline (Fig. 4B).

**Figure 4.**
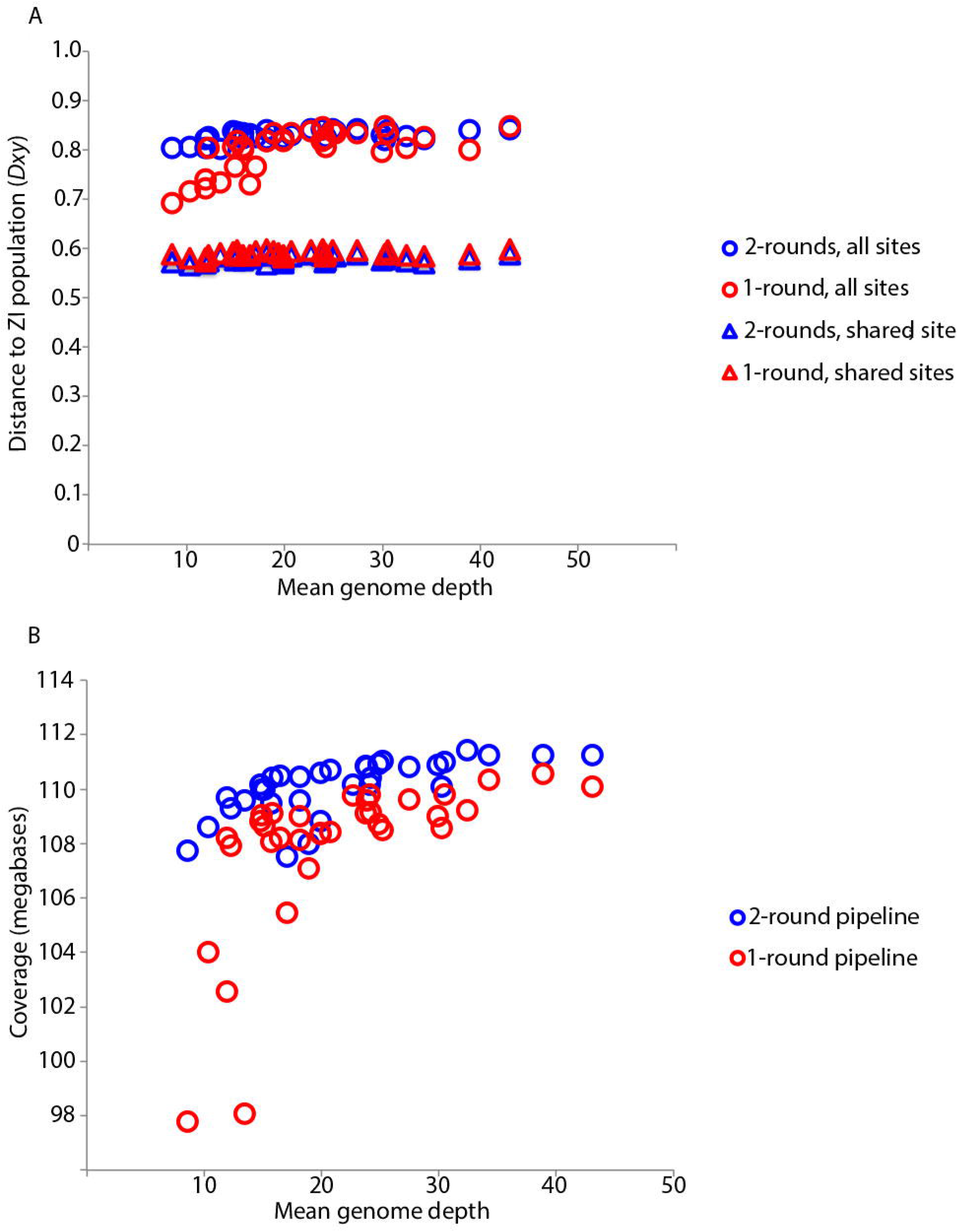
Mean sequencing depth vs genetic distance (A) from the Zambia population and depth vs coverage (B) for the POOL dataset genomes with high coverage on all chromosome arms (listed in Table S1). Circles indicate comparisons utilizing all windows with called sites, while triangles indicate comparisons including only sites called for all of the POOL and ZI genomes. Comparisons illustrate the effect of depth on genetic distance (A) and coverage (B) for genomes assembled using a single-round pipeline (red), vs our two round pipeline (blue). The two round pipeline appears to alleviate the potential downward bias present in the single-round pipeline for depths below approximately 20X, and the greater impact of depth on coverage for the single-round pipeline (B) suggests the sites added by the two round pipeline are driving the differences in distance to ZI.

### Heterozygosity

Heterozygosity can persist in fly stocks even after many generations of full-sibling mating, probably due to the presence of recessive lethal or infertile mutations, which are commonly found on wild-derived *Drosophila* chromosomes (Greenberg and Crow 1960). Especially when combined with inversion polymorphism (*e.g.* one recessive lethal is fixed on the inversion-bearing chromsomes, and a different recessive lethal is fixed on the standard arms), recombination may be unable to generate reproductively viable homozygous progeny, and residual heterozygosity may extend over much of a chromosome arm.

We report heterozygosity tracts in Table S3, including those for the Egypt EG, Kenya KM, and Zimbabwe ZK samples. The largest non-isogeneous sample in our analysis is the 205 DGRP genomes originating from Raleigh, North Carolina, USA. In spite of 20 generations of full-sib mating for the DGRP lines, considerable residual heterozygosity was maintained within the inbred lines. Overall, 12.6% of the total genomic sequence was masked due to apparent heterozygosity, and for each autosomal arm there were multiple fly lines for which the entire chromosome arm remained heterozygous. Considerably less masking was needed on the X chromosome than on the autosomes, which is expected given the increased efficacy of selection against recessive lethals and steriles in hemizygous males.

To examine the role of inversions in maintaining heterozygosity in inbred lines, we obtained inversion genotypes for each euchromatic arm of the DGRP lines from Huang *et al.* (2014). As is evident from the distribution of heterozygosity proportions for inverted vs standard autosomal arms (Fig. 5 and Table S7), more than 80% of the chromosome arms with inversion polymorphism retain over 95% heterozygosity, compared to chromosome arms lacking inversion polymorphism for which more than 80% retained less than 10% heterozygosity. These results support the role of recessive deleterious mutations residing within large inversions driving chromosome arm-wide residual heterozygosity, but fail to explain the remaining residual heterozygosity evident in the standard arm distribution shown in Fig. 5 (6.5% of non-inverted chromosome arms still retained >25% heterozygosity after 20 generations of inbreeding). While it is possible that inversion differences between sequenced and karyotyped sub-lines might exist in some cases, another explanation is that multiple recessive lethals in repulsion on a single chromosome arm might reduce the rate at which viable recombinants arise during inbreeding (Falconer 1989).

**Figure 5.**
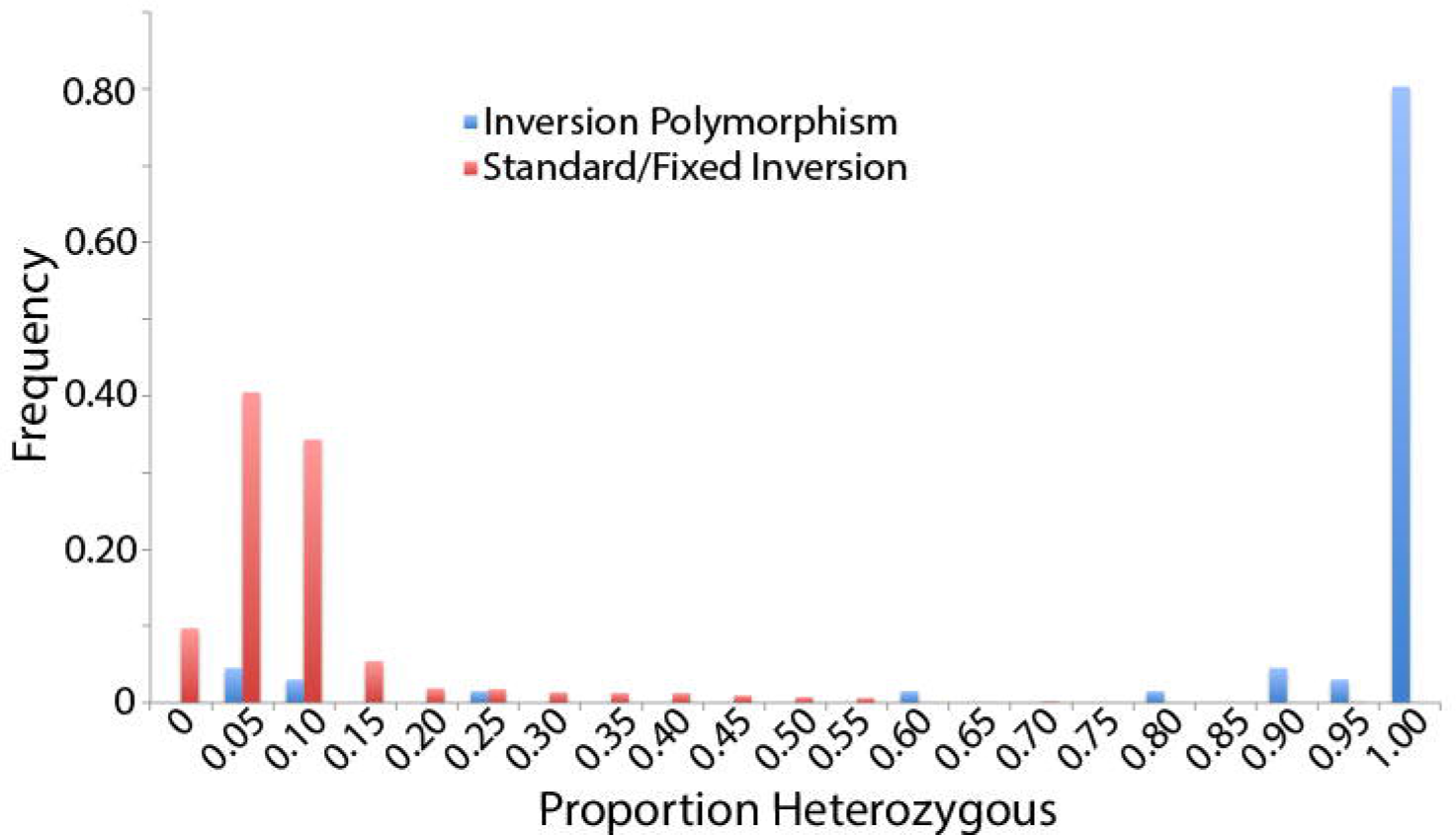
A histogram of the proportions of each autosomal chromosome arm called heterozygous from the 205 DGRP genomes. Based on the cytological analysis of Huang *et al.* (2014), red arms were reported to be free of inversion polymorphism, while blue arms contained polymorphic inversions. The greatly increased heterozygosity of the latter category illustrates the effects of inversion polymorphism on inbreeding efficacy.

In addition to true heterozygosity, artifactual “heterozygosity” (pseudoheterozygosity) can result from mismapping or other technical issues with genome assembly. For the haploid embryo genomes presented here, these positions constituted a very small proportion of total sites in a given genome (Mean = 0.00798; SD = 0.00501; Min. = 0.00025; Max. = 0.0310).

### Identity by descent

Identity-by-descent (IBD) regions passing all filters were flagged and are provided as an optional filter in the Drosophila Genome Nexus release (all IBD tracts are given in Table S6). For the DPGP2 data set, the IBD tracts we identified were essentially identical to those of Pool *et al.* (2012) and therefore are not discussed. For the AGES data set, we detected IBD for a single pair of samples from the SF South African population, but this segment included one long tract encompassing all of Chr3R and half of Chr3L. For the DPGP3 data set, we detected IBD for 20 sample pairs, constituting only 3.2% of all called bases and 0.1% of all pairwise comparisons for those 197 genomes. For the 205 DGRP genomes, IBD appeared to be more widespread, with 9.8% of all called bases flagged for masking. For a case of two IBD genomes, these base counts refer to only the masked individual, and a total of 54 IBD sample pairs were detected.

### Cosmopolitan admixture in African genomes

It has previously been noted that the introgression of cosmopolitan alleles into some African populations could have a significant influence on genetic diversity within Africa (Begun and Aquadro 1993; Capy *et al.* 2000; Kauer *et al.* 2003), and cosmopolitan admixture proportions were previously estimated for the DPGP2 data set by Pool *et al.* (2012). We repeated this analysis for all sub-Saharan genomes published here, but with improved reference panels (including more genomes, but excluding inverted arms – see Materials and Methods). All identified cosmopolitan admixture tracts are given in Table S8, and are provided as an optional sequence filter in the Drosophila Genome Nexus release.

As in the DPGP2 analysis (Pool *et al.* 2012), admixture varied considerably among populations, from <1% in the Ethiopian EM population to > 80% in the Zambian ZL population (Fig. S7). Within-population variation was also striking, as is evident from individual genome plots of admixture (Fig. 6). One important exception to the high level of inter-individual variation in cosmopolitan admixture proportions was the large DPGP3 sample (ZI) from Siavonga, Zambia. We targeted this population sample for large-scale genome sequencing for multiple reasons, including its hypothesized position within the ancestral range of *D. melanogaster* (showing maximal genetic diversity), as well as its relatively low level of cosmopolitan admixture among four genomes surveyed in the DPGP2 analysis (Pool *et al.* 2012). Our analysis of the larger DPGP3 data set illustrates that the ZI population does in fact have a very low level of cosmopolitan admixture, with the population average at 1.1% of the genome, the highest individual genome at 26%, and the second highest at 9% (Fig. 6). Looking across the genome, DPGP3 is similar to other sub-Saharan genomes in having the lowest admixture levels on the X chromosome (Fig. S8), but has a pronounced increase in the middle of arm 3R (roughly 7.6 Mb to 15.0 Mb), where up to 13 putatively admixed genomes are found in the maximal window (6.6% of the sample), compared with a genome-wide median of just 2 out of 197 individuals.

**Figure 6.**
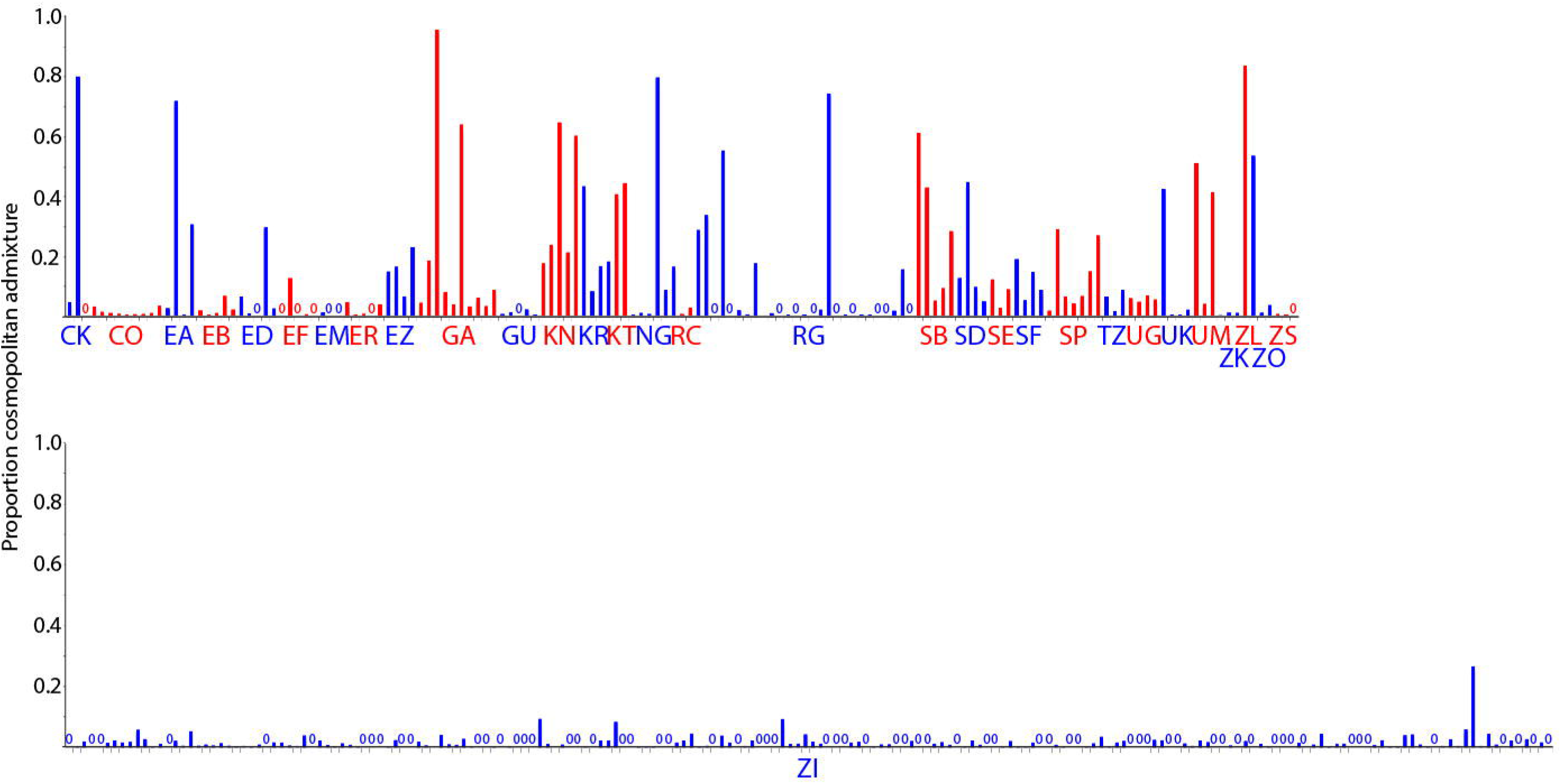
Heterogeneity in estimated cosmopolitan admixture proportions among individuals for each Sub-Saharan African population.

### Genetic diversity and structure

Although the present study is not primarily focused on population genetic analysis, we present a few simple summaries of the data to guide potential users of these assemblies. First, we estimated nucleotide diversity for all populations with multiple high coverage genomes for all chromosome arms, both including and excluding inverted arms. For the DPGP2 data set, nucleotide diversity was largely consistent with the previous estimates of Pool *et al.* (2012), although estimates for the newly assembled sequences were generally slightly higher (Table S9), perhaps due to the improved coverage of more diverse regions.

Nucleotide diversity comparisons among populations revealed similar patterns of past analyses (Table S9). The France, Egypt, and U.S. (DGRP RAL) populations had much lower diversity levels than any sub-Saharan populations (Table S9), particularly on the X. This strong reduction in diversity has been previously documented (Begun and Aquadro 1993; Baudry *et al.* 2004) and presumably results from the bottleneck that occurred during expansion out of sub-Saharan Africa. With additional sub-Saharan African genomes, as well as the expansion of the Siavonga, Zambia population to nearly 200 genomes, south-central Africa remains the most diverse portion of the *D. melanogaster* distribution. While Siavonga, Zambia still has the highest nucleotide diversity at 0.854% (Table S9), samples from Zimbabwe and inland South Africa reach 0.814% to 0.850%. The ancestral range of the species may have included much of southern Africa, unless a more recent expansion occurred with very little loss of diversity. Both eastern and western African populations were still reduced in diversity relative to southern Africa (generally 0.73% to 0.80%). Pool *et al.* (2012) reported a further, mild diversity reduction in Ethiopian highland populations was as previously described (Pool *et al.* 2012), but a lowland sample from far western Ethiopia (EA) showed little diversity reduction.

To examine the effects of inversions on diversity at the genome-scale, we estimated nucleotide diversity with inverted arms removed (Table 2). Previous analyses revealed that the effects of inversions on nucleotide diversity were not limited to regions surrounding breakpoints, but could affect entire chromosome arms (Corbett-Detig and Hartl 2012, Pool *et al.* 2012). Among the studied sub-Saharan populations, inversions appeared to have effects of both elevating and reducing arm-wide diversity (Table 2). The North American RAL sample showed less diversity elevation from inversions compared with the European FR sample.

**Table 2.** **Chromosomal arm nucleotide diversity (*π*) for populations with inversion polymorphism.** Nucleotide diversity estimates include both the total data set for a given population (Total) and excluding arms carrying inversions (Standard). “Non” denotes populations with less than two standard chromosome arms; “no inv.” denotes populations without inversion polymorphism. Values in bold indicate a difference ≥5%.

|  | X |  | 2L |  | 2R |  | 3L |  | 3R |  |
| --- | --- | --- | --- | --- | --- | --- | --- | --- | --- | --- |
| Pop | Standard | Total | Standard | Total | Standard | Total | Standard | Total | Standard | Total |
| CO | 0.0075 | 0.0075 | 0.0082 | 0.0083 | 0.0073 | no inv. | 0.0076 | no inv. | Non | 0.0058 |
| EA | 0.0071 | 0.0071 | 0.0087 | no inv. | 0.0074 | no inv. | 0.0075 | no inv. | 0.0060 | no inv. |
| EB | 0.0064 | no inv. | 0.0074 | no inv. | 0.0066 | no inv. | 0.0067 | no inv. | **0.0054** | **0.0060** |
| EG | 0.0034 | 0.0034 | Non | 0.0066 | 0.0053 | no inv. | Non | Non | Non | 0.0062 |
| FR | 0.0036 | 0.0036 | **0.0055** | **0.0061** | 0.0051 | no inv. | **0.0054** | **0.0063** | **0.0045** | **0.0058** |
| GA | 0.0077 | 0.0076 | **0.0087** | **0.0082** | 0.0077 | no inv. | 0.0080 | no inv. | 0.0066 | 0.0068 |
| GU | 0.0076 | 0.0076 | 0.0083 | 0.0084 | 0.0075 | no inv. | 0.0077 | no inv. | Non | 0.0066 |
| KN | 0.0087 | 0.0086 | **0.0061** | **0.0078** | 0.0077 | no inv. | 0.0082 | no inv. | 0.0070 | 0.0073 |
| KR | 0.0082 | 0.0085 | Non | 0.0068 | 0.0080 | 0.0081 | 0.0085 | no inv. | 0.0067 | no inv. |
| NG | 0.0076 | 0.0076 | **0.0109** | **0.0077** | 0.0075 | 0.0074 | **0.0085** | **0.0073** | **0.0054** | **0.0065** |
| RAL | 0.0041 | 0.0041 | 0.0068 | 0.0070 | 0.0062 | 0.0064 | 0.0064 | 0.0065 | 0.0052 | 0.0052 |
| RG | 0.0080 | no inv. | 0.0085 | 0.0094 | **0.0076** | **0.0081** | 0.0078 | no inv. | 0.0063 | 0.0065 |
| SB | 0.0088 | no inv. | **0.0106** | **0.0094** | 0.0082 | no inv. | 0.0083 | 0.0089 | 0.0072 | no inv. |
| SD | 0.0086 | no inv. | 0.0096 | 0.0097 | 0.0078 | 0.0078 | 0.0080 | no inv. | 0.0065 | no inv. |
| SE | 0.0083 | 0.0086 | Non | 0.0075 | 0.0079 | 0.0076 | 0.0081 | no inv. | 0.0072 | no inv. |
| SF | 0.0086 | no inv. | **0.0103** | **0.0094** | **0.0090** | **0.0085** | 0.0081 | no inv. | **0.0048** | **0.0065** |
| SP | 0.0090 | no inv. | 0.0099 | 0.0100 | 0.0083 | 0.0082 | 0.0083 | no inv. | 0.0065 | no inv. |
| TZ | Non | 0.0058 | Non | 0.0062 | 0.0077 | no inv. | 0.0075 | no inv. | **0.0056** | **0.0066** |
| UK | 0.0081 | 0.0081 | 0.0085 | 0.0086 | 0.0077 | 0.0078 | 0.0077 | no inv. | **0.0060** | **0.0065** |
| UM | 0.0080 | 0.0081 | Non | 0.0084 | 0.0078 | no inv. | 0.0068 | no inv. | **0.0075** | **0.0066** |
| ZI | 0.0089 | 0.0089 | 0.0099 | 0.0097 | 0.0083 | 0.0082 | 0.0084 | 0.0087 | 0.0076 | 0.0076 |
| ZS | **0.0089** | **0.0082** | 0.0099 | 0.0098 | 0.0083 | 0.0080 | 0.0082 | no inv. | 0.0074 | 0.0074 |

Estimation of *F*_*ST*_ and *D*_*xy*_ revealed patterns similar to those of Pool *et al.* (2012) for the DPGP2 populations (Table S9). Within the population groupings identified in that study, population differentiation was particularly low among southern African populations (mean *F*_*ST*_ = 0.0092), and somewhat elevated among Ethiopian samples (mean *F*_*ST*_ = 0.0331) – which as previously observed, showed moderate differentiation from other sub-Saharan samples (Table S9). Examination of *F*_*ST*_ restricted to standard chromosome arms indicated mainly small effects of inversions on genetic differentiation: in some cases the addition of inversions increased genetic differentiation (*e.g.* Nigeria NG vs. other sub-Saharan samples), while in other cases inversions decreased genetic differentiation (*e.g.* for comparisons involving the France or U.S. samples). Concordant with previous observations (*e.g.* Caracristi and Schlotterer 2003; Haddrill *et al.* 2005) and the hypothesized admixed origin of New World populations from European and African sources, standard arms from the North American RAL sample had consistently higher diversity than the European FR sample (Table 2), as well as closer relationships to sub-Saharan populations (Table 3).

**Table 3.** Pairwise population *F*_*ST*_ for select populations averaged across chromosome arms. Comparisons utilizing the total data set for each population is above the diagonal, and comparisons using only arms without inversions are shown below the diagonal.

|  | **FR** | **GA** | **NG** | **RAL** | **RG** | **SP** | **ZI** |
| --- | --- | --- | --- | --- | --- | --- | --- |
| **FR** | 0.0000 | 0.1898 | 0.2263 | 0.0376 | 0.2173 | 0.2213 | 0.2152 |
| **GA** | 0.2270 | 0.0000 | 0.0263 | 0.1626 | 0.0515 | 0.0955 | 0.0874 |
| **NG** | 0.2630 | 0.0133 | 0.0000 | 0.1954 | 0.0719 | 0.1128 | 0.1067 |
| **RAL** | 0.0444 | 0.1672 | 0.2000 | 0.0000 | 0.1879 | 0.1967 | 0.1911 |
| **RG** | 0.2545 | 0.0515 | 0.0631 | 0.1965 | 0.0000 | 0.0694 | 0.0595 |
| **SP** | 0.2565 | 0.0962 | 0.1034 | 0.2079 | 0.0768 | 0.0000 | 0.0130 |
| **ZI** | 0.2508 | 0.0939 | 0.1015 | 0.2025 | 0.0662 | 0.0127 | 0.0000 |

## Discussion

We have presented a set of 605 consistently aligned *D. melanogaster* genomes. Although our pipeline primarily makes use of published methods, the resulting alignments are expected to yield a better combination of accuracy and genomic coverage than standard approaches. However, the primary motivation for the Drosophila Genome Nexus is to increase the comparability of population genomic data sets, as well as make available more than 250 additional genomes, including the large Siavonga, Zambia sample.

Our effort accounts for one category of potential biases between data sets (differences in alignment methodology and data filtering), but other potential concerns should still be recognized. Although not addressed here, differences in data generation, including (but not limited to) methods of obtaining genomic DNA and sequencing platform/chemistry, may influence the resulting genomic data (Quail *et al.* 2012; Ratan *et al.* 2013; Solonenko *et al.* 2013). Our pipeline reduces the population genetic consequences of differences in sequencing depth, but depth still has an important influence on genomic coverage. Mapping success may vary according to a genome’s genetic similarity to the reference sequence, which for *D. melanogaster* is expected to have primarily cosmopolitan origin. This genetic similarity to the reference sequence will vary geographically (*e.g.* sub-Saharan genomes being more genetically distant from the reference) and across the genome (especially for admixed populations). Demography may also bias downstream population genetic analyses: for example, recent admixture and identity-by-descent are contrary to the predictions of models that assume random sampling of individuals from large randomly mating populations (thus we provide filters to reduce the effects of these specific issues).

It should be emphasized that the present DGN is primarily aimed at SNP-oriented analysis of the five major euchromatic chromosome arms. Aside from inversion-calling and the detection of short indels, we do not address the important topic of structural variation. Furthermore, the challenge of reliably aligning heterochromatin and other repetitive regions (on a population scale) awaits further technological and methodological progress.

Thorough population genetic analysis of the DPGP3 Zambia (ZI) population sample will be a topic of future analyses. However, the preliminary statistics reported here support the notion that these genomes will be widely utilize in the field of population genetics. This population continues to present the maximal genetic diversity of any *D. melanogaster* population studied to date, offering hope that it may be the least affected by losses of genetic diversity via expansion-related population bottlenecks. Unlike many sub-Saharan populations, it also contains very little cosmopolitan admixture. The availability of nearly 200 genomes from this single sub-Saharan population sample, which may have a relatively simpler demographic history than many *D. melanogaster* populations, will be an asset for studies seeking to understand the genetic, selective and demographic mechanisms that shape genomic polymorphism and divergence in large populations.

## Acknowledgements

We thank Stephen Richards for assistance with the DGRP data, J. J. Emerson for packaging the admixture HMM method, and Isaac Knoflicek for help with data management and the DGN web site. We also acknowledge the University of Wisconsin Center for High Throughput Computing (CHTC) for computational resources and assistance regarding our alignments. Funding was provided by NIH grant HG02942 to CHL, NIH grant R01 GM111797 to JEP, and support to JBL from a Ruth L. Kirschstein National Research Service Award (F32 GM106594) and from the University of Wisconsin-Madison Genome Sciences Training Program (GSTP).

## Supporting Material

**Figure S1.** Graphical depiction of the two round assembly pipeline.

**Figure S2.** Length distributions for called indels for 46 bp (blue) and 150 bp (pink) read lengths. The inset zooms in on the frequencies for lengths ≥40 bp.

**Figure S3.** Evaluation of the tradeoff between genomic coverage and error rate for the haploid caller of the Unified Genotyper; quality values ranged from 10 to 100. Resequenced genomes from the reference strain (*y*^1^ *cn*^1^ *bw*^1^ *sp*^1^) were modified to simulate realistic levels of variation. We chose a cutoff of Q75 (red) to maximize coverage and minimize error.

**Figure S4.** Evaluation of the tradeoff between genomic coverage and error rate for the diploid caller of the Unified Genotyper; quality values ranged from 10 to 100. Resequenced genomes from the reference strain (*y*^1^ *cn*^1^ *bw*^1^ *sp*^1^) were modified to simulate realistic levels of variation. We chose a cutoff of Q32 (red) to maximize coverage and minimize error.

**Figure S5.** Histogram of sequence tract lengths for sites added by our two round pipeline for the Rwandan genomes RG5 and RG33.

**Figure S6.** Number of sites added by our two round assembly pipeline in 100 kb windows across the 5 euchromatic chromosome arms for the Rwandan genomes RG5 and RG33.

**Figure S7.** Variation among African populations in estimated cosmopolitan admixture proportions.

**Figure S8.** Numbers of sub-Saharan genomes inferred to have cosmopolitan ancestry in each genomic window: (A) For the DPGP3 Zambia ZI sample, and (B) across all other sub-Saharan populations. Windows are depicted for arms X (green), 2L (blue), 2R (purple), 3L (red), and 3R (orange).

**Table S1.** Individual sequenced genomes included in this data release, including fly stock ID, genomic library ID and type/source for each library, NIH SRA access numbers, focal chromosome arms, read length, coverage and mean depth of focal chromosome arms, and the data set from which the original sequenced reads originated.

**Table S2.** Population samples from which the sequenced genomes originated. The number of sequenced individuals for each focal chromosome arm is given.

**Table S3.** Coordinates of residual heterozygosity tracts and pseudoheterozygosity tracts filtered from genomes, and proportions of true heterozygosity and total masked heterozygosity for every genome in the data set. The distinction between the masked proportion and true heterozygosity proportion is due to the presence of artefactual heterozygosity (pseudoheterozygosity) resulting from mismapping or technical issues with individual libraries.

**Table S4.** Recurrent identity-by-descent (IBD) tracts for the each data set. Only IBD tracts outside of these regions were allowed to contribute to individual totals.

**Table S5.** Inversions detected from fly stocks via cytology (DGRP; Huang *et al.* 2014) or from genomes via bioinformatics (Corbett-Detig and Hartl 2012). Note that in the case of the haploid embryo genomes, live stocks may harbor undetected inversion polymorphism. “INV/ST” indicates known polymorphism. “INV/?” indicates that inverted reads were detected, but the genome was heterozygous in this region. Blank cells indicate inversions that were untested or unreported for this genome/stock.

**Table S6.** Regions of IBD masked from the analyzed genomes, including both individual genomes identified for each tract. See the methods for a detailed description of IBD detection and filtering criteria, and Table S4 for the excluded recurrent IBD regions.

**Table S7.** Inversion polymorphism and proportion heterozygosity on each focal chromosome arm for each DGRP genome, illustrating the role of inversions in maintaining heterozygosity in spite of considerable inbreeding effort. “Pseudoheterozygosity” corresponds to the proportion of a chromosome arm prior to normalization, and “Corrected heterozygosity” corresponds to the proportion of a chromosome arm following normalization.

**Table S8.** Regions of cosmopolitan admixture masked in Sub-Saharan African genomes.

**Table S9.** Genome-wide genetic differentiation and nucleotide diversity for populations with multiple high-coverage focal chromosomes, averaged across the five focal chromosome arms. Values below the diagonal are *F*_*ST*_, values above the diagonal are *D*_*xy*_, and bold values on the diagonal are nucleotide diversity. Distance from the *D. melanogaster* reference genome is given in the bottom row.

**Table S10.** Individual chromosome arm measures of genetic differentiation and nucleotide diversity for populations with at least two high-coverage sequences. Values below the diagonal are *F*_*ST*_, values above the diagonal are *D*_*xy*_, and bold values on the diagonal are nucleotide diversity. Distance (*D*_*xy*_) from the *D. melanogaster* reference genome is given in the bottom row.

